# Western diet feeding since adolescence impairs functional hyperemia probed by functional ultrasound imaging at adulthood and middle age: rescue by a balanced ω-3:ω-6 polyunsaturated fatty acids ratio in the diet

**DOI:** 10.1101/2023.12.21.572595

**Authors:** Haleh Soleimanzad, Clémentine Morisset, Mireia Montaner, Frédéric Pain, Christophe Magnan, Mickaël Tanter, Hirac Gurden

## Abstract

Obesity is a devastating worldwide metabolic disease, with the highest prevalence in children and adolescence. Obesity impacts neuronal function but the fate of functional hyperemia, a vital mechanism making possible cerebral blood supply to active brain areas, is unknown in organisms fed a high caloric Western Diet (WD) since adolescence. We mapped changes in cerebral blood volume (CBV) in the somatosensory cortex in response to whiskers stimulation in adolescent, adult and middle-aged mice fed a WD since adolescence. To this aim, we used non-invasive and high-resolution functional ultrasound imaging (fUS). Functional hyperemia is compromised as early as 3 weeks of WD and remains impaired thereafter. Starting WD in adult mice does not trigger the profound impairment in sensory-evoked CBV observed in young mice, suggesting a cerebrovascular vulnerability to WD during adolescence. A balanced ω-6:ω-3 polyunsaturated fatty acids ratio in WD achieved by docosahexaenoic acid supplementation is efficient to restore glucose homeostasis and functional hyperemia in adults.

## Introduction

According to the World Health Organization, obesity is particularly affecting the youngest populations. Very low in the 1970’s, the prevalence of overweight and obesity among the 5-19 years old have reached 18% in 2016^1^. This range is by far the one with the highest increase for indicators of obesity^2^. Obesity is a global health threat resulting from excessive consumption of energy-dense food and insufficient calorie expenditure^3^. In consequence, obesity is primarily defined by the excessive fat accumulation in the body^4^. This disease predisposes patients to numerous pathologies such as type 2 diabetes (T2D), nonalcoholic fatty liver disease and cardiovascular diseases^3^ by hampering the function of organs such as the pancreas, the liver and the heart^5^. Obesity also impacts several aspects of brain function, including sensory and cognitive processing, in rodent models^6,7^ and in humans^8,9^. In obese patients, very high intake of energy is due to the frequent ingestion of Western Diet (WD) enriched in both saturated fatty acids and refined carbohydrates. WD is notably characterized by a deleterious unbalanced ω-3:ω-6 polyunsaturated fatty acids (PUFAs) ratio and causes gluco- and lipo-toxicity in cerebral tissues^10,11^. This impacts *in fine* neuronal and cerebrovascular function^12,13^.

The very high energy demand of neuronal activity is fulfilled by a vascular supply which has to be instantly regulated^14^. In brain networks, coordinated neurovascular coupling mechanisms make possible functional hyperemia, i.e. the increase in cerebral blood flow (CBF) and volume (CBV) in response to increased synaptic activity^15^. Failure in proper regulation of functional hyperemia is observed in diseases such as chronic cerebral hypoperfusion^16^ leading to vascular dementia and pathogenesis underlying Alzheimer’s disease^17^. Obesity is also associated with reduced functional hyperemia in adults and elderlies^12^ and, together with T2D^18^, is a metabolic condition where the brain is susceptible to pathologies like ischemic stroke and vascular dementia^16^. Much effort has been made to characterize brain cerebrovascular activity in metabolic diseases in adult and old ages^19^ and in the context of ischemia (in rodents^20^ and humans^13^) but changes in functional hyperemia from adolescence to mid age in lean and obese organisms have not been examined.

To follow up functional hyperemia, several complementary imaging techniques were used in animals^21^ and humans^22^. Thanks to the typical 50 fold increase in terms of sensitivity to CBF obtained using ultrafast Doppler imaging^23,24^, a new concept of ultrasound neuroimaging emerged during the last decade^25^. Such functional Ultrasound (fUS) imaging brought significant insights into the understanding of brain function with neurodevelopmental, sensory, motor and cognitive studies^26^. Recent fUS imaging also demonstrated its ability to assess brain dysfunction in diseased models^27^ but metabolic pathologies were not examined using this efficient technique for intact *in vivo* imaging recordings so far.

We decided to follow-up changes of energy metabolism and sensory-evoked functional hyperemia in adolescent animals chronically fed with WD. Our data demonstrate that WD resulted in obese and prediabetic mice characterized by hyperglycemia, glucose intolerance, insulin resistance and hyperinsulinemia compared to lean mice. These obese mice presented an early and sustained impairment in functional hyperemia from adolescence to mid age. We further showed that using docosahexaenoic acid-supplemented WD to balance the ω-3:ω-6 PUFAs ratio during adolescence prevents the deleterious effects on metabolic and cerebrovascular functions linked to WD in adult obese mice.

## RESULTS

### Starting WD at adolescence induces profound metabolic deficiency in adolescent, adult and middle-aged mice

First, we compared main metabolic functions of WD-fed mice (5317 kcal/kg), starting at adolescence and maintained under WD until middle age, to age-matched regular diet (RD)-fed control mice (2791kcal/kg). We characterized the metabolic phenotype of 1 week (W), 2W, 3W, 2 months (M), and 10M WD-fed and RD-fed groups (Fig. 1a). Although no significant differences of body weight nor body composition were shown after 1 or 2 weeks of WD (1W and 2W groups, Fig. 1b,c), 3 weeks of WD caused significant body weight (3W group, Fig. 1b) and fat mass (Fig. 1c) gain in WD-fed mice. Mice with 3 weeks of WD displayed a significant rise in fasting blood glucose (Fig. 1d) compared to RD mice, a major metabolic defect that is not present at 1 or 2W of WD. We observed that this deleterious effect of WD on glycemia persisted until adulthood (2M group) and middle age (10M group). In addition, Oral Glucose Tolerance Test (OGTT) showed increased glucose intolerance (Fig. 1e) and Insulin Tolerance Test (ITT) showed decreased insulin sensitivity (Fig. 1f) which worsened from 3 weeks to 10 months of WD. We also observed increased basal insulinemia in all WD-fed groups (Fig 1g-k), with a loss of glucose-stimulated insulin secretion. Thus, feeding mice with WD at adolescence led to obese and prediabetic states in adolescence, adulthood, and mid-age. WD-induced obesity in mice is an accurate model to mimic the metabolic syndrome in humans^28^.

**Fig. 1.**
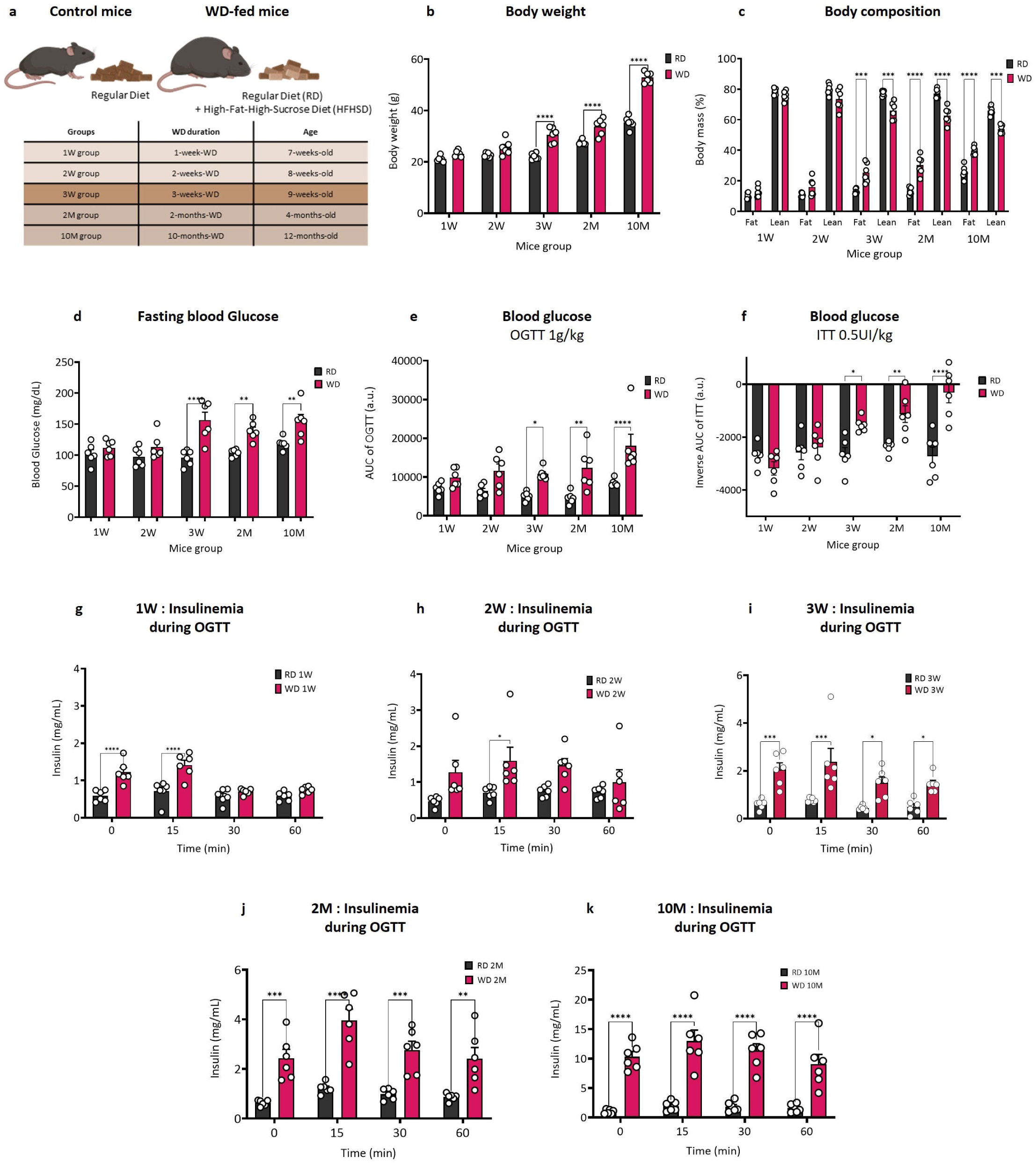
Glucose metabolism in mice fed a WD at adolescence is altered from 3 weeks to 10 months of diet compared to age-matched RD mice. **(a)** Experimental groups. Abbreviations indicate the duration of diet. For example, 1W corresponds to 1 week of WD. Since diet starts at 6 weeks of age (adolescence), 1W group corresponds to 7 week-aged mice. Note that the WD leading to obesity is a mixture of RD and high-fat high-sucrose diet. **(b-i)** Metabolic phenotyping of obese mice fed a WD for a duration of 1, 2, 3 weeks or 2, 10 months (1W, 2W, 3W, 2M and 10M groups) and their respective age-matched control mice groups fed with RD. **(b)** Body weight, **(c)** fat and lean mass and **(d)** overnight fasting blood glucose. **(e)** Area under the curve (AUC) of blood glucose during oral glucose tolerance tests (OGTT) and **(f)** inverse AUC of blood glucose during insulin tolerance tests (ITT). **(g-i),** Plasma insulin levels from blood samples taken during the OGTT test in **d**. Note that the threshold of metabolic dysfunction starts at 3 weeks of WD. Metabolic data are represented as mean ± SEM and were acquired on the same mice for each diet group at a given age, n=6 per diet group at a given age. P values were calculated using two-way ANOVA with post hoc Bonferroni tests. *P<0.05, **P<0.01, ***P<0.001, and ****P<0.0001. a.u., arbitrary units; ns, not significant.

### fUS imaging reveals rapid and long-lasting impairment of functional hyperemia in mice fed a WD since adolescence

We imaged sensory-evoked hemodynamic responses non-invasively in the barrel cortex of anesthetized WD-fed and age-matched RD-fed mice in response to mechanical whisker stimulation at 2Hz. We observed transcranial cerebrovascular responses using fUS imaging with 15MHz ultrasonic array placed over a coronal plane covering the somatosensory cortex (Fig. 2a). To detect activated brain areas during whisker stimulation, we computed Z-scores based on general linear model analyses (GLM) with Bonferroni correction for multiple comparison (normalization by the total number of pixels, see Methods). Computing the fUS activation map clearly showed an area of activation in the contralateral barrel cortex in response to whisker stimulation in all mice groups (Fig. 2b and 2d). To further characterize the amplitude of sensory evoked-CBV, we quantified the mean variation of relative CBV (mean ΔrCBV, % of baseline, Fig 2c left and 2f) over the responding activated pixels (ΔrCBV map, Fig. 2e). We observed that highly activated pixels were further located inside the left barrel cortex^25^ in response to right whiskers mechanical stimulation. Then, we computed the averaged rCBV time-course through stimulation trials (Fig. 2c right and 2f) and observed that it systematically showed a transient and stimulus-locked increase in activated areas in all groups. We also calculated the mean ΔrCBV amplitude (at the plateau between 10 to 30 seconds of whisker stimulation) over baseline (mean %ΔrCBV, Fig. 2g). No significant differences were observed by paired comparison of WD and RD groups with 1 week, 2 weeks and 10 months of diet but we could reveal a significant reduction in ΔrCBV in groups with 3 weeks and 2 months of WD. Thus, feeding mice with WD resulted in strong and long-term impairment of functional hyperemia in the somatosensory cortex of mice.

**Fig. 2.**
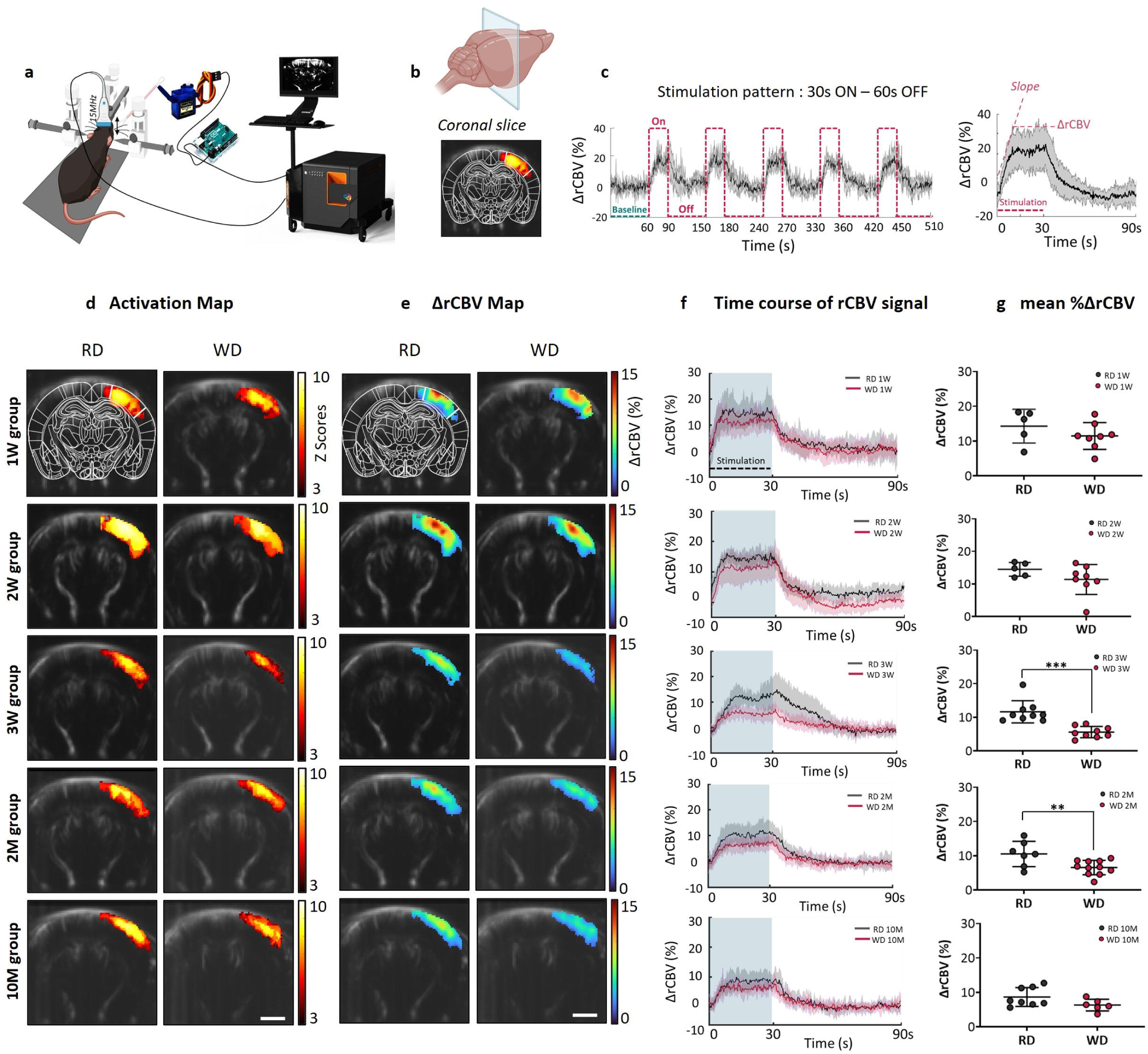
Impaired functional hyperemia to whisker stimulations in adolescent, adult and middle-aged mice fed a WD since adolescence compared to age-matched RD mice. **(a)** Experimental setup. The intact skull and intact skin of the head-fixed anesthetized mouse is insonified in a coronal plane continuously with a 15 MHz central frequency ultrasound probe during mechanical whisker stimulation in the anesthetized mouse. **(b)** fUS imaging of the somatosensory-induced activity at Bregma −1.7 mm with superimposition of the activated pixels (color scale) over the brain vasculature (grey scale). Mean activation map is showing significantly activated pixels in the left barrel cortex area following stimulation of right whiskers with a custom-made mechanical stimulator. Background map centered on the barrel cortex coordinates^76^. **(c)** Time course changes of relative CBV (rCBV) signal. Left: time course changes of rCBV as a percentage of baseline in the barrel cortex (ΔrCBV(%)). The total recording time for a single trial was 510 s, with a 60 s baseline, a 30 s stimulation and a 60 s recovery time. 5 consecutive whisker stimulation trials were acquired (curves in transparency defines Standard Deviation, n = 11 RD mice). Right: mean time course changes of ΔrCBV(%) averaged over 5 trials (mean ± SD, n = 11, from **(c)**). Rise slope of the fUS signal and ΔrCBV at the maximum plateau of the vascular response are locked to whisker-stimulation onset. **(d)** Activation map. Same representation than **(b)** for all diet duration groups. **(e)** ΔrCBV map. Mean variation of relative CBV (ΔrCBV) values are represented over responding pixels in response to five stimulation trials. Color intensity goes from dark blue (no change during stimulation) to red (strong CBV increase during the 30 s stimulation). Scale bar in **(a-b)** 2 mm. **(f)** Time course of rCBV signal. Whisker stimulation-evoked variation in ΔrCBV is plotted as a percentage change of baseline (%ΔrCBV). Note the plateau of maximum %ΔrCBV between 10 s and 30 s of whisker stimulation **(g)** Mean %ΔrCBV averaged at the maximum plateau between 10 s and 30 s stimulation from **(f)**. 1W group: 14.3 ± 4.8% for RD (n=5) vs 11.5 ± 3.9% for WD-fed mice (n=8, p=0.26); 2W group: 13.3 ± 3.8% for RD (n=5) vs 11.4 ± 4.6% for WD-fed mice (n=8, p=0.45); 3W group: 11.7 ± 3.3% for RD (n=9) vs 5.6 ± 1.7% for WD-fed mice (n=9, p=0.0002); 2M group: 10.5 ± 3.7% for RD (n=7) vs 6.5 ± 2.1% for WD-fed mice (n=11, p = 0.01) and 10M group: 8.7 ± 2.7% for RD (n=8) vs 6.3±1.7% WD-fed mice (n=6, p=0.09). Note that vascular dysfunction starts at 3 weeks of WD.

### Sensory-evoked functional hyperemia dynamics during development and aging are different in lean versus obese mice

We further analyzed fUS signals and computed values of rise slope and maximum ΔrCBV variation from the signal time course (Fig. 2c) at different ages (Fig 3a-b). RD mice showed a significant decrease in the slope value in the 3W compared to 1W group (Fig. 3a). This decrease was still present at 2 months and 10 months of diet. Interestingly, this result fits with the physiological decline of cerebral hemodynamics during adolescent brain development in humans^29^. The same pattern of slope decrease was present in WD-fed mice as they age but the decline was further amplified during obesity (Fig 3a). Interestingly, ΔrCBV at middle age in the 10 moths RD-fed mice was significantly decreased compared to young 1-2 week control groups (Fig. 3b). This explains the similar decreased hemodynamic response magnitude in WD and RD observed in the 10M group (compare Fig. 2g and Fig. 3b). In fact, aging dynamics of ΔrCBV is very different between obese and control mice (Fig. 3b-c): a sudden decrease of the ΔrCBV amplitude occurred in 3W WD-fed mice and remained impaired in 2M, whereas in RD-fed animals the decrease was present at middle age only (10M)^7^. This observation is in favor of the existence of premature aging mechanisms that could occur in WD-fed mice^7,30^.

**Fig. 3.**
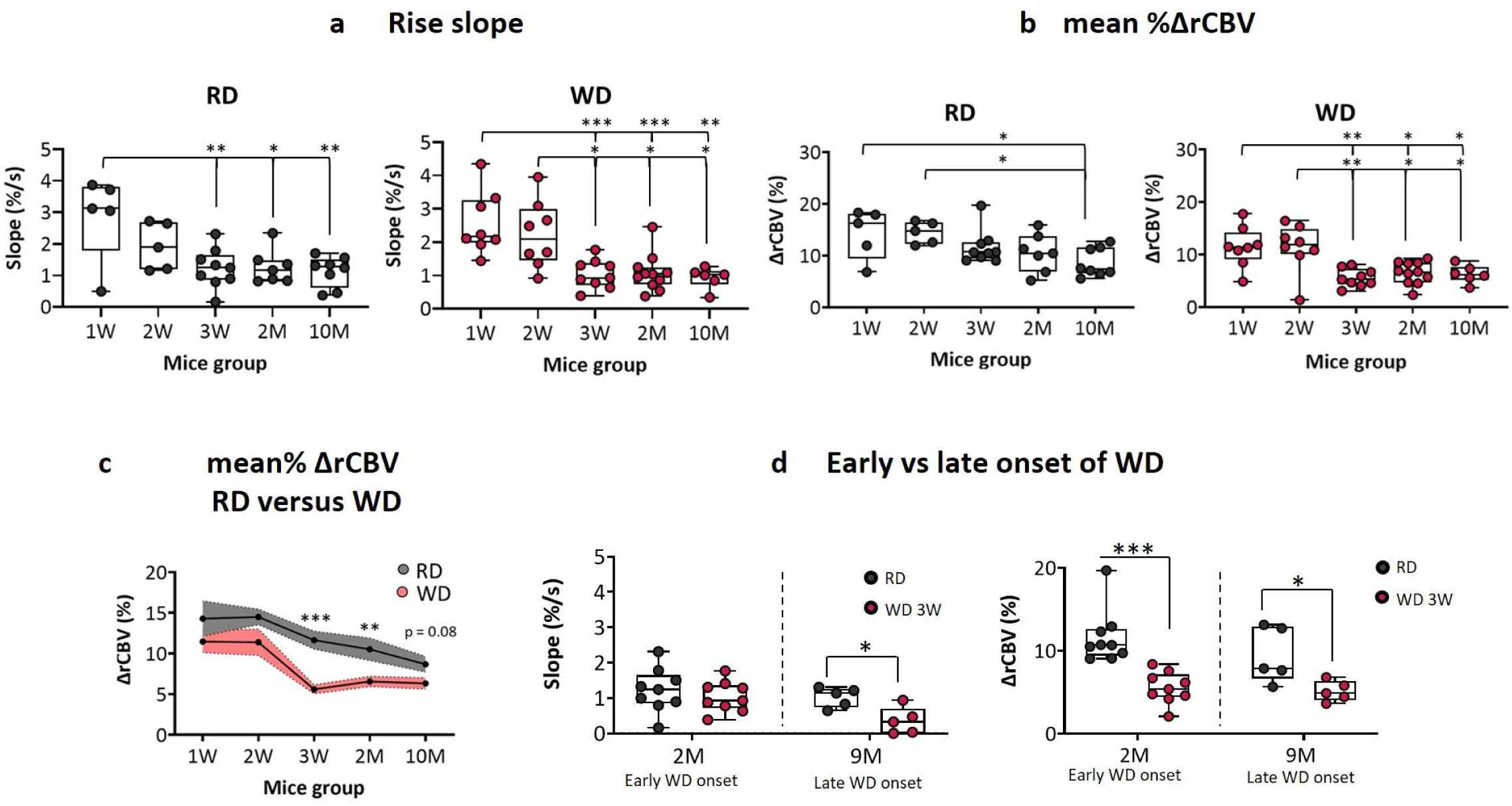
Functional hyperemia dynamics during mouse development are different in lean and obese mice. **(a)** Aging effect on the rise slope of CBV time course represented in fig 2g for RD groups (left panel) and WD groups (right panel). The rise slopes of the CBV time course signal were calculated on the first 5 s of sensory stimulation. RD mice: 2.8 ± 0.48% of CBV increase per second in 1W group (n=5), 1.9 ± 0.22 in 2W group (n=5), 1.2 ± 0.14 in 3W group (n=9), 1.2 ± 0.2 in 2M group (n=7) and 1.1 ± 0.13 for 10M group (n=8). For WD-fed mice: 2.5 ± 0.31% of CBV increase per second in 1W group (n=8), 2.2 ± 0.2 in 2W group (n=8), 1.0 ± 0.16 in 3W group (n=9), 1.0 ± 0.13 in 2M group (n=11) and 0.9 ± 0.18 in 10M group (n=6). **(b)** Aging effect on the mean CBV calculated from time course in fig 2g for RD groups (left panel) and WD groups (right panel). Data points are the same than in fig 2h. **(c)** Comparison of RD and WD groups at different ages. Data points are the same than in **(b)**. **(d)** Beginning WD at adolescence not adulthood triggers profound impairment in functional hyperemia. Left: Right: Results in **(a-g)** are presented as mean ± SD for 1W (n = 5 for RD mice, n = 8 for WD mice), 2W (n = 5 RD, n = 8 WD), 3W (n = 9 RD, n = 9 WD), 2M (n = 7 RD, n = 11 WD) and for 10M group (n = 8 RD, n = 6 WD). Mean P values were calculated using ANOVA for intragroup aging effects with posthoc Bonferroni’s multiple comparisons test and unpaired t-test to compare RD vs WD at different ages. *P<0.05, **P<0.01, ***P<0.001, and ****P<0.0001.

### Beginning WD in adolescence is more deleterious for functional hyperemia than in adulthood

Since the deleterious effect of WD on the hippocampal function in juvenile rodents was shown to be stronger than in adults^6^, we wondered whether the age of WD onset could be an important factor for the negative impact of western diet on CBV. To answer this question, adolescent versus adult mice were WD-fed during 3 weeks, resulting respectively in 2 month-aged animals (2M, early WD onset, Fig. 3d-left panel) and 9 month-aged mice (late WD onset, Fig. 3d-right panel). Starting WD at adulthood resulted in a slightly impaired slope and ΔrCBV values 3 weeks after WD onset (Fig. 3d right and left panel) whereas it strongly compromised cerebrovascular activity in adolescent mice (Fig. 3d-right panel). This result indicates that WD onset is a crucial factor in triggering altered functional hyperemia and further underlies the greater vulnerability of the adolescent brain to WD toxicity compared to adult brains^31^.

### Using docosahexaenoic acid (DHA) supplementation to balance the ω-3:ω-6 PUFAs ratio to 1 restores glucose tolerance and insulin sensitivity in WD-fed mice

WD in humans is characterized by an unbalanced ω-3: ω-6 PUFAs ratio (from 1/10 to 1/20), in favor of ω-6 PUFAs^32^. The WD used in our experiments has a ratio of 1/11, in the range of WD consumed by humans. Since DHA, a ω-3 PUFA highly concentrated in the brain, has promising protective effects on the cardiovascular system^33^, including on the cerebral blood vessels in ischemic models^34^, we decided to add DHA to WD to restore an accurate ω-3:ω-6 PUFAs ratio of 1. To evaluate how this balanced ratio of fatty acids affects metabolism, adolescent mice were fed with either RD, WD or WD enriched with DHA (WD+DHA) during 2 months before reaching adulthood. Body weight (Fig. 4a) and fat mass (Fig. 4b) were significantly increased in both WD-fed and WD+DHA-fed groups compared to RD. Interestingly, fasting glycemia in WD+DHA mice had an intermediate profile, significantly reduced compared to WD but higher compared to RD (Fig 4c). In addition, supplementing mice with DHA resulted in selective recovery of glucose tolerance (AUC of the OGTT, Fig. d) and insulin sensitivity (AUC of the ITT, Fig. e) in the WD+DHA compared to the WD group (Fig. 4d-e). DHA supplementation also resulted in decreased baseline levels of insulinemia (t0) and decreased glucose-induced insulinemia (t15, Fig. 4f) in the WD+DHA group compared to WD mice, further showing the gain of efficacy in insulin action under balanced ω-3:ω-6 PUFAs ratio^35^. We conclude that restoring a balanced ω-3:ω-6 PUFAs by DHA supplementation restores major metabolic parameters in obese mice.

**Fig. 4.**
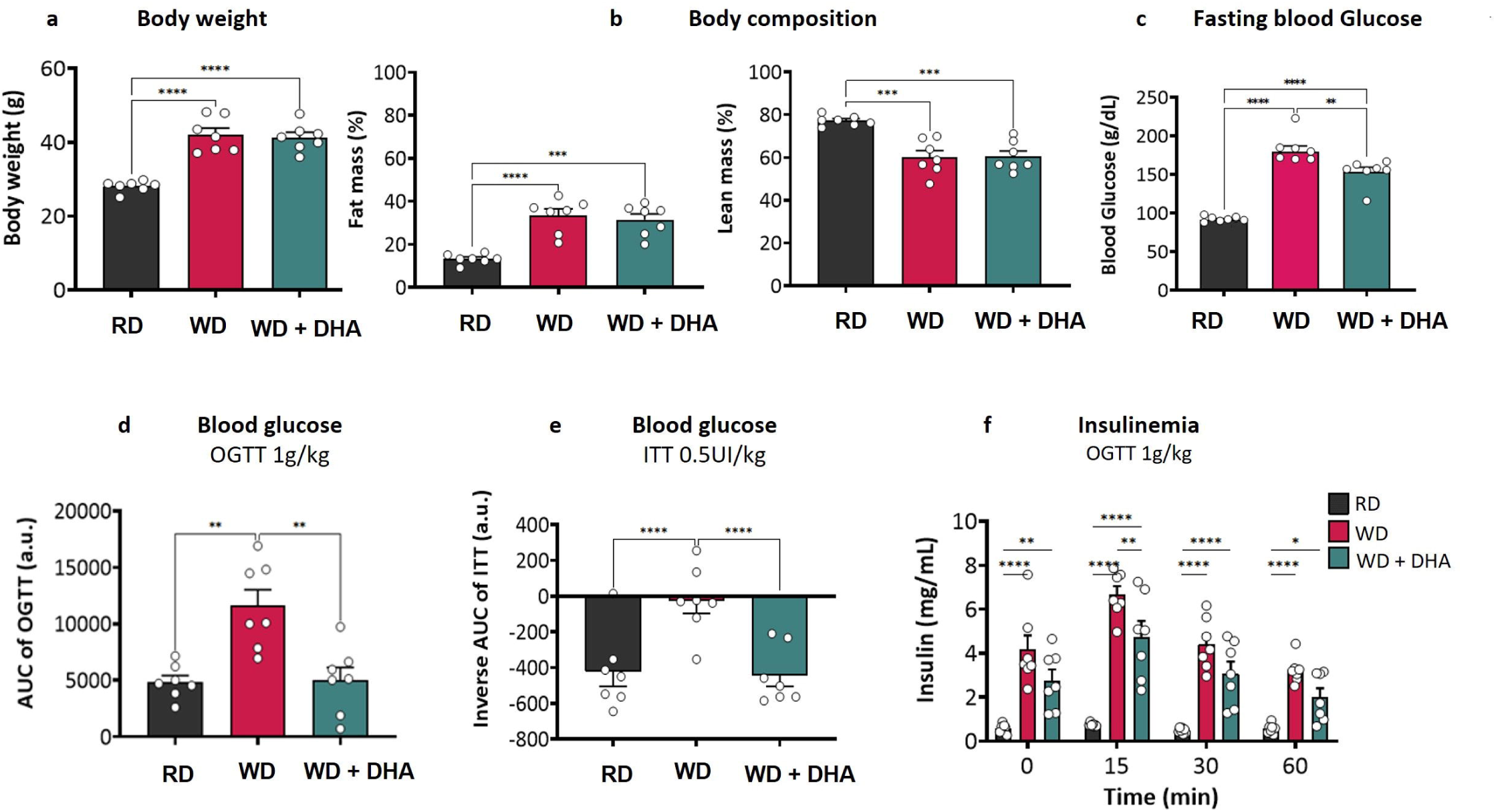
Impairment in glucose metabolism in obese mice due to WD is canceled by feeding with WD balanced in the ω-3:ω-6 PUFAs ratio. Metabolic phenotyping of adult RD-fed versus adult obese mice fed either WD or WD+DHA for a duration of 2 months starting at adolescence. **(a)** Body weight, **(b)** fat and lean mass, **(c)** overnight fasting blood glucose, **(d)** AUC of blood glucose during OGTT and **(e)** inverse AUC of blood glucose during insulin ITT. **(f)** Plasma insulin levels from blood samples taken during the OGTT test in **(d)**. **(g-i)** same analysis and representation for fUS imaging than Fig.2. c-d. Data are represented as mean ± SEM and were acquired on the same mice from each diet group, n=7 per diet group. P values were calculated using two-way ANOVA with post hoc Bonferroni tests. *P<0.05, **P<0.01, ***P<0.001, and ****P<0.0001. a.u., arbitrary units; ns, not significant.

### A ω-3:ω-6 PUFAs ratio of 1 prevents functional hyperemia deficits in WD-fed mice

To test whether the metabolic improvement triggered by balanced ω-3:ω-6 PUFAs PUFAs ratio could positively influence functional hyperemia, we recorded whisker-evoked CBV increase by fUS imaging. As observed before, the mean activation map (Fig. 5a) and the mean ΔrCBV map (Fig. 5b) showed an intense activation across the barrel cortex in RD-fed mice whereas the activated area was restricted in WD-fed mice. Interestingly, RD and WD+DHA mice showed a similar signal increase (Fig. 5a-b). We also observed a similar temporal evolution of the CBV signal in WD+DHA and RD-fed mice (Fig. 5c). These results indicate that DHA supplementation cancels the decrease in fUS signal observed with WD alone.

**Fig. 5.**
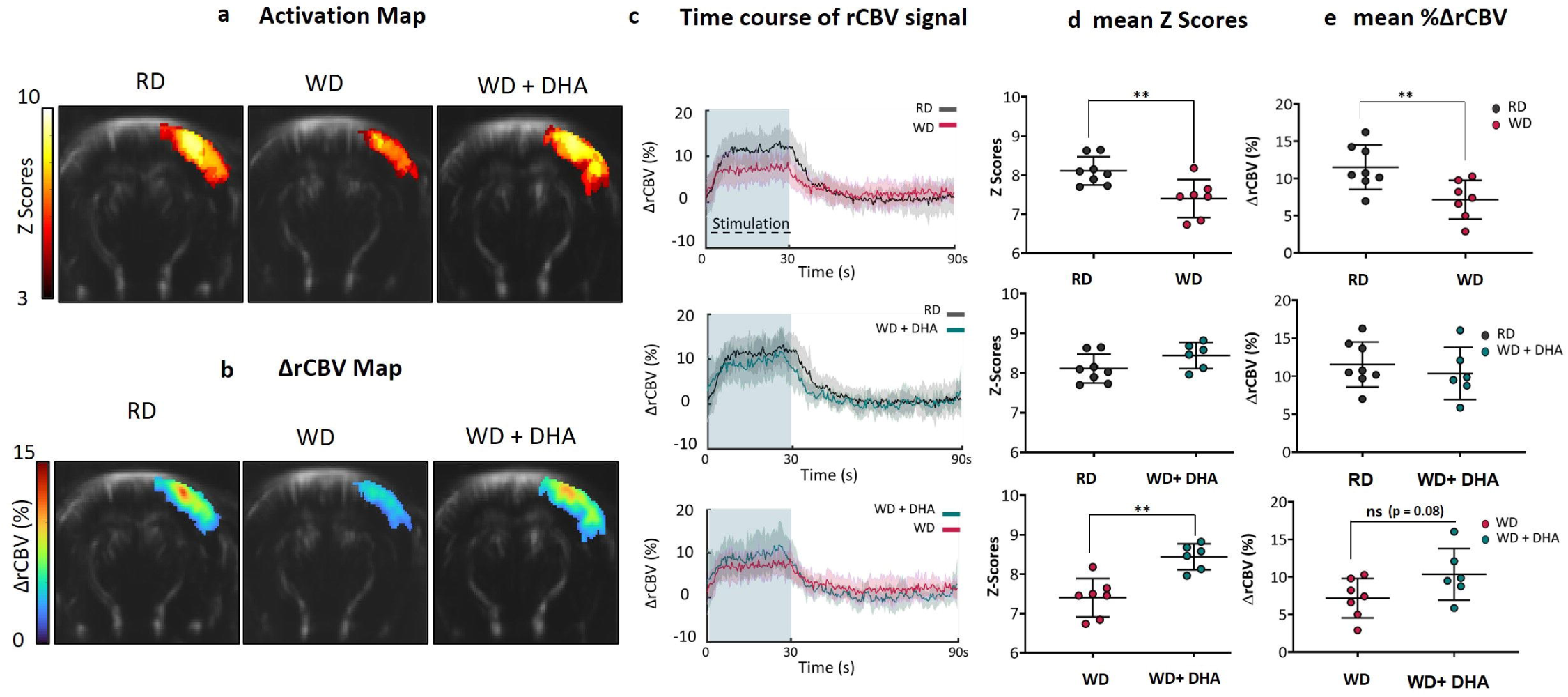
The amplitude of functional hyperemia is restored in adult WD-fed mice with a balanced ω-3:ω-6 PUFAs ratio. **(a)** Activation map. Same representation that Fig. 2d for RD, WD and WD+DHA groups. **(b)** ΔrCBV map. Same representation than Fig. 2e for RD, WD and WD+DHA groups. **(c)** Avergaged time course changes of relative CBV as a percentage of baseline in the barrel cortex (ΔrCBV(%)) for RD, WD and WD+DHA groups. **(d)** Number of the activated pixels recorded in the activation map (quantified as Z-scores). Mean Z-score is 8.11 ± 0.36 in RD-fed mice (n = 8), 7.40 ± 0.48 in WD-fed mice (n=7) and 8.44 ± 0.33 in WD+DHA-fed mice (n=6). **(e)** Mean %ΔrCBV at maximum plateau (10-30 s of whisker stimulation). Mean values are 11.56 ± 2.9 for RD-fed mice (n=8), 7.2 ± 2.6 for WD-fed mice (n=7) and 10.37 ± 3.4 for WD+DHA-fed mice (n=6). Data are presented as mean ± SD. P values are calculated using unpaired t-test. P<0.05, **P<0.01, ***P<0.001, and ****P<0.0001.

Computing the average Z-scores of all significant pixels (Fig. 5d) from the ΔrCBV map (Fig. 5b) revealed a decreased activated area in WD compared to RD (Fig. 5d, top), an equal mean between WD and WD+DHA mice (Fig. 5d, middle) and a significantly higher value for WD+DHA group compared to WD (Fig 5d, bottom). This shows again the protective role of DHA on functional hyperemia in obese mice. In addition, the WD-fed mice showed a significant decrease in ΔrCBV compared to RD controls (Fig. 5e, top) while WD+DHA and RD mice were not different (Fig. 5e, middle). WD+DHA mice presented a trend towards ΔrCBV increase compared to WD-fed mice but this difference was not significant (Fig. 5e, bottom). Thus, a balanced ω-3:ω-6 PUFAs ratio resulted in the improvement of sensory-evoked cerebrovascular responses compared to an unbalanced ratio. This result complements and strengthens, in the context of adolescence obesity, previous observations that DHA is beneficial to brain vascular function in obesity^36^.

## Discussion

In this study, feeding WD mice at adolescence resulted in glucose intolerant, insulin resistant and hyperinsulinemic obese prediabetic mice. This metabolic disorder starts three weeks after WD onset at adolescence and is amplified at adulthood and middle age. fUS imaging recordings demonstrate an early deficiency in functional hyperemia in mice after the beginning of WD at adolescence. fUS further revealed decreased functional hyperemia at middle age in lean mice and a long-lasting cerebrovascular impairment at adulthood and middle age in obese mice. A balanced ω-3:ω-6 PUFAs ratio prevents functional hyperemia deficits induced by WD at adulthood.

Childhood and adolescence obesity is a worldwide devastating disease and has become a major global public-health concern^37^. It is a major drive of type 2 diabetes risk^38,39^ at early ages. In line with previous studies^40,41^, our diet-induced obesity model revealed disrupted metabolic regulation with early impaired insulin sensitivity and decreased capacity in glucose regulation in the beginning of WD. Our murine model is also characterized by an increased fasted glycemia, which defines the appearance of a prediabetic phenotype in mice and fits with the adolescent metabolic syndrome. At this early age, failure rates in achieving optimal glycemic control are high and young diabetic patients rapidly develop severe complications, including multiple defects in brain function^8^. The cerebrovascular activity is highly sensitive to pathological metabolic changes in the organism. Hampered functional hyperemia in our obese prediabetic mice is detected as soon as 3 weeks after WD onset and these mice never recover. This fits the human cerebrovascular picture. Increased BMI in adults is detrimental for CBF regulation^12,42^ Young obese adults in their early twenties present lower CBF than healthy young subjects^43^. Importantly, obesity-induced insulin resistance in children and adolescents is the most important risk factor of vascular diseases in adulthood^44^. In fact, early-onset T2D is a more aggressive disease from a vascular standpoint than adult T2D. In that sense, diabetes in obese adolescent patients further decreases CBF compared to obese adolescents^45^.

Indeed, our data show that the decline in function hyperemia is not only present at adolescence but is amplified and sustained in older, adult, and middle-aged mice fed a WD since adolescence. Previous publications described an early impairment in functional hyperemia with WD starting at adulthood^46,47^. Here, we further revealed that the same duration of WD is more harmful for metabolic and cerebrovascular regulation in adolescent mice compared to adult mice. This pattern indicates that onset time of WD is a crucial factor in triggering altered long-term functional hyperemia and highlights the greater vulnerability of the adolescent brain to WD toxicity. Our results are in line with the cognitive impairment triggered by exposure to obesogenic diet in adolescent^6,48^ but not in adult mice^6,31^. They complete studies reporting global vulnerability of the neuronal and vascular network of the adolescent brain facing chronic neurotoxicity (substance abuse, mild traumatic brain injury, stress, WD…)^31,49,50^.

Very interestingly, we also revealed two periods of normative functional hyperemia decrease in lean, metabolically healthy mice. First, decreased cerebrovascular activity in lean adolescent mice fits with the physiological decrease of cerebral hemodynamics during brain development in healthy human adolescents^29,51,52^. Secondly, impaired functional hyperemia in middle-aged mice is likely to correspond to normative decrease of cerebrovascular activity in middle-aged humans^53^. More interestingly, our results about the blunted functional hyperemia in young adolescent obese mice, together with publications showing faster decreased CBF in the olfactory bulb of obese mice^7^ and the accelerated vascular aging among adolescents with obesity and/or T2D^30,54^, suggest a premature aging of the cerebrovascular function in young obese organisms.

Insulin resistance is pivotal to the metabolic syndrome: all body and metabolic markers show significant correlation to insulin resistance in human adolescents^55^. It is associated with impairment in cardio- and cerebrovascular function^5,45,56^. In our experiment, insulin resistance is detected very early, 3 weeks after the beginning of WD, which fits with previous publications reporting very fast impact of hypercaloric diet on insulin sensitivity^41,57^. This pattern is consistent with the deep impairment of cerebrovascular function in Zucker diabetic and insulin resistant rats^58^. Interestingly, a neutracetical approach with enriched DHA not only improves beta cell function in periphery but decreases insulin resistance in preclinic models and in human adolescents^35,59^ and our own results are in favor of a recovery of insulin sensitivity by DHA supplementation resulting in a balanced ω-3:ω-6 PUFAs ratio.

Personalized nutrition for the prevention of T2D are among the new tools to pediatricians and pediatric diabetologists^60^. A higher intake of ω-3 PUFAs may be beneficial in the management of obesity^61,62^, for cardiac and brain vascular function^36^, especially in adolescents^63^. Long-chain ω-3 PUFAs fulfil crucial structural and functional roles in the body, notably in the human brain^64^, and low ω-6:ω-3 PUFAs ratio is important for proper brain development in young ages^32^. In contrast, ω-6 PUFAs are pro-inflammatory^65^, with higher intakes being linked to increased incidence of obesity and metabolic diseases^66^. One efficient way to restore a balanced ω-3:ω-6 PUFAs ratio is to use DHA supplementation in the WD and new nutritional methods include the use of ω-3 enriched diets^67^ as complementary approaches to pharmacological treatments against obesity and obesity-related health risks. Using this nutritional approach, we show an improvement of all metabolic parameters as well as sensory-evoked CBV responses in obese mice. Our data complement and strength in the context of adolescent obesity previous observations in adults that ω-3-PUFA-enriched diets^68,69^, and particularly DHA supplementation, is beneficial to brain vascular function^34,36,63^. Obviously, similarly to adult obesity, preventing weight gain during child development is the best strategy to avoid any overweight-related disorders. However, once overweight is settled in adolescents, increasing ω-3 PUFAs contents in the diet could be a first beneficial approach for cerebrovascular protection. Of note, in our experimental conditions, metabolic improvement induced by a balanced ω-3:ω-6 PUFAs ratio was not accompanied by weight loss. Thus, in our model, DHA is not likely to influence peripheral lipotoxicity directly but rather to contribute to local protective mechanisms in the brain^70^.

In conclusion, thanks to the high spatiotemporal resolution of fUS, we could efficiently sample functional hyperemia in intact mice at different ages. fUS made it possible to i) show a mild decrease in functional hyperemia in adolescent and middle-aged lean mice as previously shown in murine models^71^ and clinics^72^ ii) demonstrate for the first time an early impairment of sensory-evoked CBV response in adolescent obese and prediabetic mice similar to what was described in several brain pathologies^17^. Further work is necessary to pinpoint the molecular/cellular sources of deficient functional hyperemia in mice fed a WD since adolescence. Still, the beneficial neutraceutical approach using balanced ω-3:ω-6 PUFAs ratio by DHA supplementation to restore functional hyperemia could serve as an efficient strategy to decrease WD toxicity on metabolic homeostasis and cerebrovascular activity.

## Acknowledgements

H.S. has a Postdoctoral Fellowship award from NIH and AXA. M.M. has a doctoral Fellowship award from Idex Université Paris Cité. This work was supported by the AXA Research Fund. We thank the animal core facility “Buffon” of the University of Paris Cité / Jacques Monod Institute for animal care.

## Author contributions

H.S and C.M equally contributed to the work. They designed the experiments, acquired and analyzed the imaging data along with M.M. for metabolic phenotyping. F.P. and C.M. edited the article. M.T. and H.G. equally contributed to the work. They designed the experiments and wrote the article together with H.S, C.M., M.M., F.P. and C.M.. M.T. and H.G. supervised the project.

## Competing financial interests

The authors declare no competing financial interests.

## Methods

All animal experiments were approved by the committee for animal care of Université Paris Cité and by the French Ministry of Research (agreement #17629) in accordance with the European directive 2010/63/UE.

### Experimental groups

Male C57BL6/J mice were hosted in animal housing starting at 5 weeks of age. After one week of habituation with all mice under regular diet (RD, Safe A04 diet, 2791kCal/kg, 10% fat, 3,8% sugar), i.e. at six weeks of age (in the rodent peri-adolescence^73^), half were shifted to a mixed RD and high fat-high sucrose diet (Western Diet (WD), Safe HF230 diet, 5317 kcal/kg, 60% fat, 27% sugar). To follow the aging effect on WD mice and their control age-matched RD mice, metabolic phenotyping and imaging were performed in 5 groups of animals at 1, 2 and 3 weeks of WD, then 2 months (adult mice) and 10 months (middle-aged mice) of diet. This means that the age of the animals in these groups was respectively 7, 8 and 9 weeks then 3,5 (adult) and 11,5 months (middle age). For the sake of clarity, we used the terms 1W, 2W, 3W, 2M, 10M for all mice to rather indicate to the WD duration (Fig. 1a). In order to study the effects of a balanced ω-3: ω-6 polyunsaturated fatty acids ratio (PUFAs, n-6:n-3=1), a WD diet supplemented with docosahexaenoic acid (DHA) was provided to mice starting at 6 weeks of age (WD +DHA group). For the WD used here, 5% DHA is needed to restore a proper ω-3:ω-6 ratio in PUFAs content of the WD (Safe diet).

### Metabolic Phenotyping

A total of 51 C57BL/6J male mice were used for metabolic phenotyping of all groups. Mice underwent Echo MRI body mass composition (EchoMRI-900 Whole Body Composition Analyzers, Echo Medical Systems, Houston, TX). Body weight was measured weekly all along the study. Animals were accustomed to daily manipulation. Fasting blood glucose was assessed after an overnight fasting via the tail vein using Glucofix® Tech (A.Menarini diagnostics, Rungis, France). Oral glucose tolerance tests (OGTT) were performed after 5 hours of fasting to evaluate the impact of WD on glycemia. Mice received glucose (1g/kg of body weight) by oral gavage and glycemia was monitored using Glucofix® Tech during 120 min. Plasma insulin levels were quantified using a Christal Chem ELISA assay from blood samples taken during the test. Insulin tolerance tests (ITT) were performed after 5 hours of fasting (0.5 UI/ kg of body weight, Novo-Nordisk, La Défense, France) and glycemia was measured until 45 min, following the same protocol as OGTT.

### Animal preparation for functional US (fUS) imaging

fUS acquisitions were performed on a total of 106 C57BL/6J male mice. Animals were anesthetized via intramuscular injection of a ketamine chlorydrate (80 mg.kg-1) and xylazine (16 mg.kg-1) cocktail. The anesthesia level was adjusted, if necessary, throughout the experiment. The body temperature was controlled by a rectal probe and maintained at 37 ± 0.5 °C by a feedback-controlled heating pad.

### Whisker Stimulation

Since the power Doppler signal is proportional to CBV^74^, we could measure neuronal activation in the barrel cortex during whisker stimulation through neurovascular coupling. An acquisition consists of a 60s baseline followed by five successive trials. Each trial consists of 30s of whisker stimulation at 2Hz, followed by 60s of recovery (no stimulation). A custom-made mechanical whisker stimulator was built using a servomotor controlled by an Arduino Uno microcontroller interfaced with the imaging system through TTL signal to synchronize the stimulation pattern with imaging sequence.

### Ultrasound Sequence

After anesthesia, the animal was placed on a stereotaxic frame to stabilize its head. Ultrasound imaging acquisitions were performed through the intact skin and skull with a 15-MHz ultrasound probe (128 elements, 15 MHz central frequency, 100 x 100 μm2 spatial resolution, Vermon, Tours, France). The probe was placed upon the brain with acoustic coupling gel to image the coronal plane at Bregma – 1.70 mm comprising the barrel cortex. The transducer was connected to an ultrafast ultrasound scanner (Iconeus One, Iconeus, France / Inserm Accelerator of Technological Research, Paris, France). Data were acquired by emitting continuously groups of 11 plane waves tilted from −10° to +10° at 5.5 kHz pulse repetition frequency. The backscattered echoes of each group were summed to get compound images at 500 Hz framerate that allows correct sampling of the blood signal. Doppler images were computed from blocks of 200 compound images averaged after appropriate filtering (see “Data Processing” section). fUS programming of custom transmit/receive ultrasound sequences was done in Matlab (version R2021b, MathWorks, USA), using software-based architecture of the scanner.

### Data Processing of fUS

Noise and tissue motion were discarded from the Doppler signal with a spatiotemporal clutter filter based on the singular value decomposition (SVD) of the ultrasound raw data. After removing the signal associated with the 60 first singular values, the time fluctuations of the remaining Doppler signal were incoherently averaged to get the final Power Doppler signal, proportional to the cerebral blood volume (CBV). Finally, a single Power Doppler image was computed from 200 compound images every 400 ms. Each pixel of the final Power Doppler images is 100 x 100 μm2 in plane. The slice thickness is 300 μm. Filtered Power Doppler images of each animal were registered to a reference mouse and linearly detrended in the spatial and temporal domains. For each mouse, we computed the relative CBV (rCBV) time course, to get rid of the inter-individual variability of cerebral perfusion. We defined the rCBV as the percentage change from the initial 60 s baseline.

### Activation Maps

For each animal, activation maps were computed with a single-subject general linear model (GLM) approach as described elsewhere^75^. Data analysis was performed using Matlab (version R2021b, MathWorks, USA). The design matrix used to detect cortical activation specific to the whisker stimulation only included the stimulus pattern as a regression signal. Statistical parametric maps were generated for each session, including a z-map, a p-value map, a stimulation map, and a baseline map. Individual delta cerebral blood volume (ΔCBV) maps were obtained by dividing the stimulation matrix by the baseline matrix. To determine which pixels achieved statistical significance considering the comparison of all the pixels in the z-map, we considered p-value < 0.05 statistically significant and further performed Bonferroni correction for multiple tests comparison. The pixels meeting the defined criteria were considered activated. For each animal, an activation map displaying the activated pixels’ z-score and a delta CBV map displaying the delta CBV of the same pixels were produced and overlaid onto the corresponding gray scale mean Doppler image. Mean activation maps were obtained after registration and averaging of these activation maps within groups.

### Functional Hemodynamic Response

For each animal, the CBV time course was computed from a 7*7 pixels box centered on the maximal z-score pixel of the activation map, after applying a median filter to the z-map. The pixel times-series of the box were averaged spatially and temporally to get one mean trial per session (i.e. per animal). Then, mean trials were averaged within groups for comparison between WD and age-matched controls at different stages and statistical analysis. The final rCBV values for statistical analysis correspond to the mean percentage change of the last 20 s of stimulation (meaning at the maximum plateau from 10 to 30s of stimulation) to the initial 60 s baseline.

### Statistics

Results of metabolic phenotyping, whisker-based texture discrimination task, multiplex cytokine/chemokine assay on plasma are expressed as the Mean ± standard error of the mean. After the validation of normal distribution by a Shapiro-Wilk test, statistical analysis for intergroup comparison was performed in GraphPad Prism (software version 9.1.2) using the following tests: metabolic phenotyping, two-way ANOVA (for the first set of experiments) or one-way ANOVA (for the second set dealing with DHA) followed by post-hoc Bonferroni test for two-by-two comparisons. Differences between groups are considered significant when the probability level is less than 5% (p-value<0.05), except for multiple comparisons where p-value was adjusted as required. In the figures, the degrees of significance are indicated by * p-value<0.05; ** p-value<0.01; *** p-value<0.001; **** p-value<0.0001.

## Notes

### Competing Interest Statement

The authors have declared no competing interest.

